# Testing the genomic link between intraspecific mating traits and interspecific mating barriers

**DOI:** 10.1101/2023.08.14.553230

**Authors:** Leeban H Yusuf, Sonia Pascoal, Peter A Moran, Nathan W Bailey

**Affiliations:** School of Biology, University of St Andrews, St Andrews, Fife KY16 9TH, UK; Department of Haematology, University of Cambridge, Cambridge, United Kingdom; A-LIFE, Section Ecology & Evolution, Vrije Universiteit Amsterdam, Amsterdam, The Netherlands

**Author notes:** **Corresponding author** (LHY), (NWB).

**Keywords:** barrier loci, genotype-phenotype, reproductive isolation, sexual selection, speciation genomics

## Abstract

Differences in interspecific mating traits such as male sexual signals and female preferences often evolve quickly as initial barriers to gene flow between nascent lineages, and they may also strengthen such barriers during secondary contact via reinforcement. However, it is an open question whether loci contributing to intraspecific variation in sexual traits are co-opted during the formation and strengthening of mating barriers between species. To test this, we used a population genomics approach in natural populations of Australian cricket sister species that overlap in a contact zone: *Teleogryllus oceanicus* and *Teleogryllus commodus.* First, we identified loci associated with intraspecific variation in *T. oceanicus* mating signals, advertisement song and cuticular hydrocarbon (CHC) pheromones. We then separately identified candidate interspecific barrier loci between the species. Genes showing elevated allelic divergence between species were enriched for neurological functions, indicating potential behavioural rewiring. Only two CHC-associated genes overlapped with these interspecific barrier loci, and intraspecific CHC loci showed signatures of being under strong selective constraint between species. In contrast, 10 intraspecific song-associated genes showed high genetic differentiation between *T. commodus* and *T. oceanicus* and two had signals of high genomic divergence. Significant increased differentiation in sympatry supported a history of asymmetrical reinforcement driven primarily by divergence in sympatric *T. commodus* populations. The overall lack of shared loci in intra vs. inter-specific comparisons of mating trait and barrier loci is consistent with limited co-option of the genetic architecture of interspecific mating signals during establishment and maintenance of reproductive isolation.

## Introduction

Determining whether and how sexual selection leads to speciation is a longstanding goal of speciation genomics, but also one of the most experimentally difficult (Ritchie, 2007; Servedio & Boughman, 2017; Mendelson & Safran, 2021). Evaluation of this idea requires identifying whether loci that cause intraspecific differences in secondary sexual traits also contribute to the origin, elaboration, or maintenance of interspecific reproductive barriers (Wilkinson *et al*., 2015). Such information is necessary to separate any signal of sexual selection during speciation from that caused by natural selection or neutral evolution, both of which can also act on sexual traits (Servedio & Boughman, 2017). Distinguishing distinct outcomes attributable to sexual selection is also complicated by the fact that the contribution of sexual selection to overall reproductive isolation between diverging populations may vary over time and space. Although theoretical and empirical studies have described ecological conditions favouring speciation via sexual selection (Kirkpatrick & Ravigné, 2002; Ritchie, 2007; Servedio & Boughman, 2017), few studies have identified loci underlying sexual traits and examined the forces acting on them during species divergence.

An effective way to study the link between the genetic basis of reproductive isolation and sexual traits is to test whether sexually selected loci overlap with candidate barrier loci (Brand *et al*., 2020; Turbek *et al*., 2021), but this approach is often hindered by the difficulties of identifying loci associated with sexual traits, identifying barrier loci between species, and then disentangling sources of selection that have historically acted on each (Wilkinson *et al*., 2015; Ravinet *et al*., 2017). Sexual selection on mating phenotypes may also play distinct roles during different stages of speciation, influenced partly by spatial context (Kulmuni *et al*., 2020). For example, interspecific differences in mating traits can result in mismatches between heterospecific signalers and receivers and cause reproductive isolation to be generated during primary divergence and reinforced later on in the speciation process (Coyne and Orr, 2004; Ritchie, 2007). However, overlapping sets of predictions make it difficult to disentangle genomic selection underlying initial build-up reproductive isolation versus that underlying reinforcement during secondary contact (Ortiz-Barrientos *et al*., 2004; Servedio *et al*., 2011; Merrill *et al*., 2012; Garner *et al*., 2018).

Figure 1 illustrates an experimental approach to determine the statistical association of loci implicated in intrasexual selection with those implicated in interspecific mating barriers at different stages of speciation, and then separate overlapping predictions caused by the dual role of sexual trait divergence as a cause and consequence of speciation. The approach combines intraspecific genotype-phenotype association tests with interspecific genome scans for divergent selection, the latter capitalizing on combined effects of local selection and gene flow – which generate identifiable heterogeneity in genomic landscapes – to search for barriers to gene flow (Ravinet *et al*., 2017). Regions underlying sexual trait differences that are also implicated in reproductive isolation are expected to show heightened genetic differentiation compared to the genomic background (Wilkinson *et al*., 2015). An important distinction can be made to identify loci implicated in sexual differences arising from reinforcement to avoid maladaptive hybridisation during secondary contact: these are predicted to show enhanced genetic differentiation in sympatry compared to allopatry (Garner *et al*., 2018).

**Figure 1.**
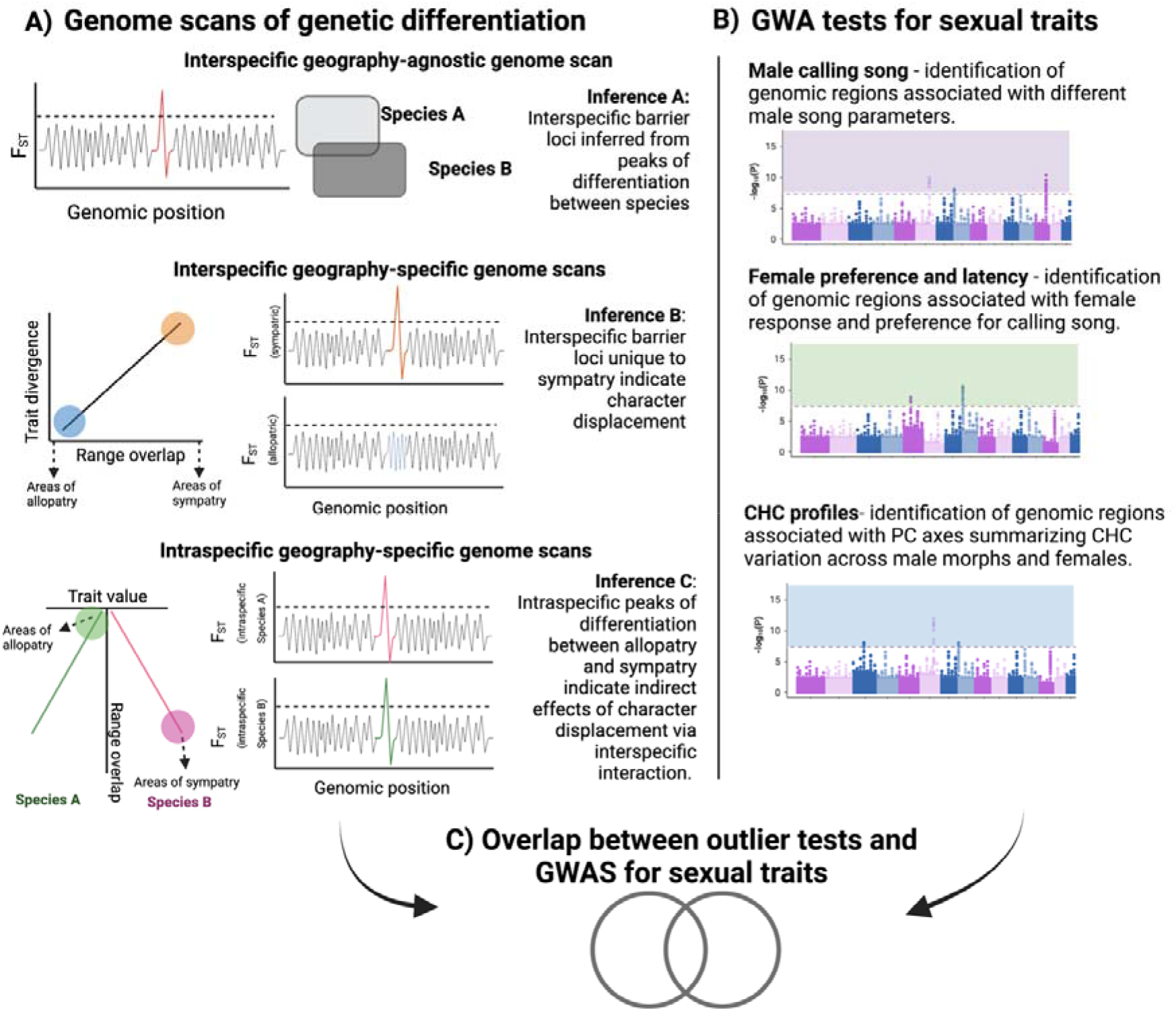
Combinatorial approach for detecting sexually selected loci implicated in reproductive isolation between *T. oceanicus* and *T. commodus*. **A)** Genome outlier scans for barrier loci using genetic differentiation in three comparisons: between species only (top row), between species in different spatial contexts (middle row), and within species in different spatial contexts (bottom row). Contrasts between loci identified in each enable inference about genes implicated in species divergence versus reproductive character displacement. **B)** Genome-wide association tests for sexual traits within species. **C)** Overlapping genomic regions found in both approaches may be promising candidate premating barrier loci.

Compared with genomic investigations of ecological differentiation (Seehausen *et al*., 2014), few causal loci underlying mating phenotypes between species have been identified. Nevertheless, those studies that have detected candidate loci using genotype-phenotype associations have most often identified oligogenic architectures for mating signals and preference between incipient species (Ding *et al*., 2016; Merrill *et al*., 2019; Brand *et al*., 2020; Turbek *et al*., 2021; Unbehend *et al*., 2021). Consistent with such genetic architecture, there is evidence that signals and preferences can co-evolve by genetic coupling, for example via pleiotropy or close physical linkage (Hoy *et al*., 1977; Butlin & Ritchie, 1989; Boake, 1991; Shaw & Lesnick, 2009; Xu & Shaw, 2019, 2021). However, assortative mating caused by phenotype matching unrelated to signal-preference coevolution, and the maintenance of gametic phase linkage disequilibrium through assortative mating, can complicate efforts to identify loci underlying genetically-coupled signal-preference architectures (Dieckmann & Doebeli, 1999; Kopp *et al*., 2017).

In this study, we used genotype-phenotype association tests and population genomic analyses across a geographical transect containing a contact zone of Australian field cricket sister taxa (*Teleoryllus* spp.) to test the genomic link between intraspecific mating traits and interspecific barrier loci. *T. oceanicus* and *T. commodus* are a classic system for studying acoustic sexual signalling and do not show evidence of ecological differentiation or habitat preference despite contact in sympatric regions of their Australian ranges (Otte & Alexander, 1983). They differ in long-range calling song used for mate location and female choice (Hill et al. 1972; Bailey and Macleod 2014), and recent behavioural analyses suggest discrimination by both sexes resulting in robust, symmetrical behavioural isolation between species (Moran *et al*., 2018). Though the species readily hybridise in the laboratory, population genetic analysis along a 2,500 km latitudinal transect of their Australian range showed no evidence for recent hybridisation in the wild (Moran *et al*., 2018). Using two extensive genomic datasets (Moran *et al*., 2018; Pascoal *et al*., 2020) and a new, high-quality chromosome-contiguous genome assembly for *T. oceanicus* (Zhang et al. in review), we characterised the genetic architecture of male calling song and CHC variation and identified loci associated with patterns of divergent selection across 16 sympatric and allopatric populations. Elevated differentiation in sympatry and inferred historical gene flow provided evidence of reinforcement. Loci related to intraspecific variation in chemical pheromones and male advertisement song features were localised throughout the genome, but the two signal types showed no co-localisation. Song loci were more likely to overlap with putative interspecific barrier loci than loci associated with chemical pheromones. The findings imply limited co-option of loci underlying intraspecific mating traits during formation or maintenance of interspecific mating barriers. However, those loci that are co-opted are likely biased toward sexual trait modalities that are less constrained by ecological factors; such loci may therefore play a prominent role in speciation.

## Methods and Materials

### Sampling, read mapping and filtering

To characterize patterns of genetic variation across the genome and understand the speciation histories of *T. commodus* and *T. oceanicus*, we used genomic data from 8 allopatric populations of *T. commodus* in southern Australia, and 4 populations of *T. oceanicus* in northern Australia that are allopatric to *T. oceanicus* (Figure 2A) (Moran *et al*., 2018)(Supplementary Table 1). We also used genomic data from individuals of both species sampled within 4 sympatric populations spanning a contact zone in the middle of the geographical transect. In northern Queensland, we also obtained samples for one additional contact zone population containing *T. oceanicus* and a third Australian species in the genus, *T. marini.* In total, raw, demultiplexed, reduced representation sequence data (RAD-seq) for 16 populations of allopatric and sympatric *T. oceanicus*, *T. commodus* and *T. marini* individuals of both sexes were used. These raw reads were previously published by Moran *et al*. (2018).

**Figure 2:**
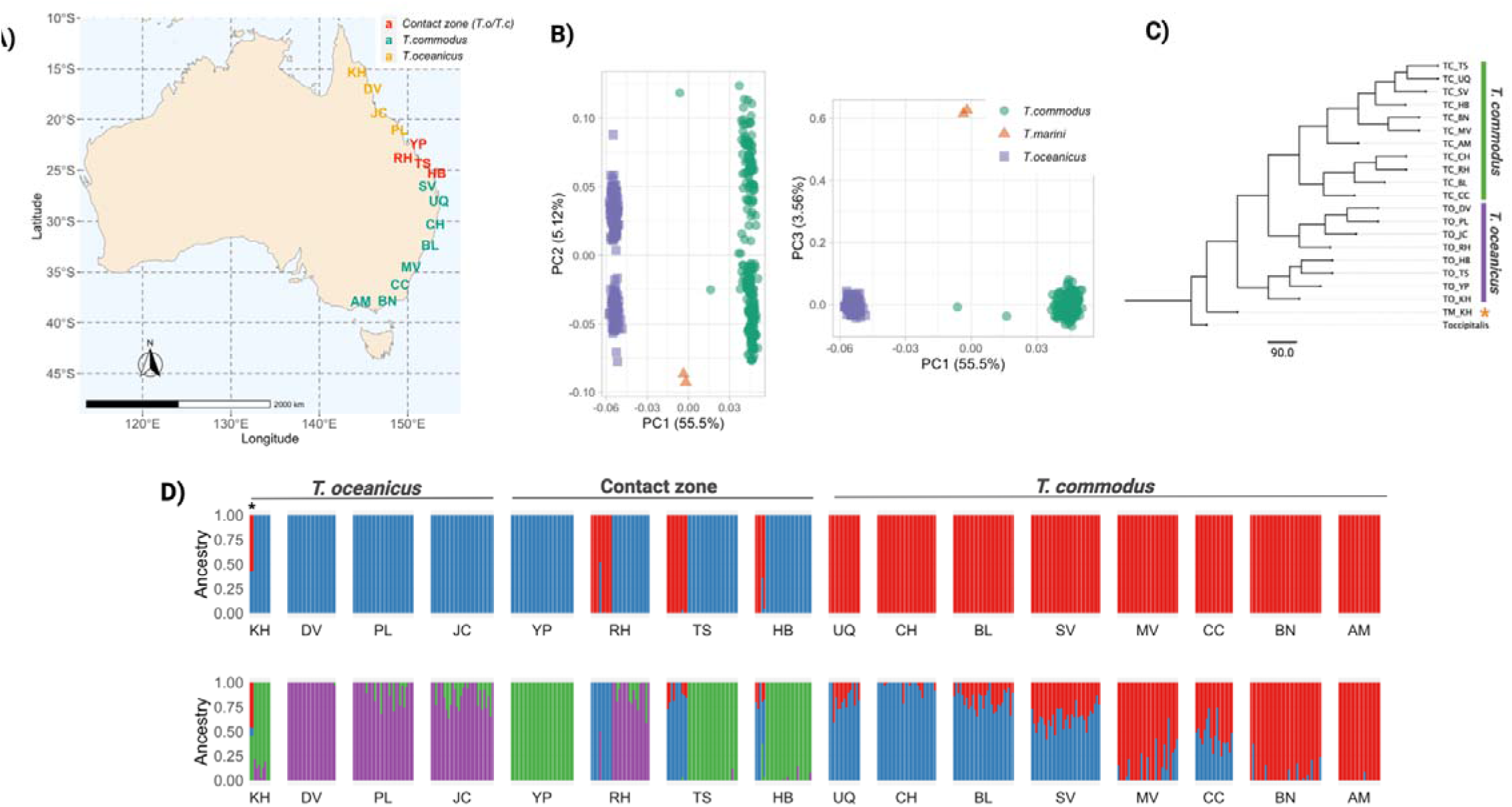
Geographic sampling scheme and population structure. **A)** T. oceanicus, T. commodus and T. marini population sampled in Moran, et al (2018) and used in this study. Colour denote allopatric and sympatric populations of *T. commodus* and *T. oceanicus*. Asterisks denote population (KH) where *T. marini* was found. **B)** PCAs of genetic variation across all samples an 4,163 linkage-pruned SNPs. **C)** Multispecies coalescent tree of populations and species**. D)** Inferred admixture fractions for allopatric and sympatric populations under K=2 (top panel) and K= (bottom panel). Asterisk denotes *T. marini* samples, which shows mixed ancestry.

Reads were trimmed using fastp v.0.20.1 (Chen et al., 2018) with default parameters, then mapped to a *T. oceanicus* reference genome (Zhang et al. in review) using BWA mem (Li, 2013; Md *et al*., 2019). We used samtools (v. 1.11) (Li et al., 2009) to index, sort, mark and remove duplicate reads from each sample and Picard (Institute, 2019) to add read groups to each sample. Variant calling was performed using bcftools mpileup and call function with the following parameters: -C 50/ -m -f GQ,GP. Variants were filtered for minimum depth (10), max missingness (0.5), minimum quality score (30), minor allele count (3) and individuals with missing genotypes at half or more of all variable sites were removed. In total, 1,451,414 variants were called and 89,045 were retained following filtering. Additionally, after removing individuals with more than 50% missing sites, we retained 417 out of 465 individuals for analysis.

### Population structure and phylogenetic reconstruction

We used PLINK (v1.90b) to assess population structure for all retained samples. To assess only independent, non-linked genetic variation, we further filtered our dataset of 89,045 biallelic SNPs down to 4,163 using the following parameters in PLINK: ‘--indep-pairwise 50 10 0.01’ to remove variants with r^2^ larger than 0.01 every 10bp in 50kb windows. We reconstructed phylogenetic relationships between allopatric and sympatric populations of *T. commodus*, *T. oceanicus* and *T. marini* using *T. occipitalis*, an Asian congener, as an outgroup. This involved mapping whole-genome sequence data of *T. occipitalis* from Kataoka *et al*. (2020) to our reference *T. oceanicus* genome using BWA mem (Li, 2013; Md *et al*., 2019) as above. We used samtools (v. 1.11) (Li et al., 2009) to index, sort, mark and remove duplicate reads. Variants were called and filtered for minimum depth (10) and minimum quality score (30). We merged filtered variants of *T. occipitalis* and *T. oceanicus*, *T. commodus* and *T. marini* to produce a comprehensive SNP dataset for gene flow inference, phylogenetic inference and with this inferred a species tree using multispecies coalescent inference and the aforementioned phylogenetic SNP dataset via SVDquartets and PAUP (v. 4a)(Chifman & Kubatko, 2014). To minimize confusion we call the first variant set of 89,045 variants our ‘population genomic variant set’, and the latter variant set containing our outgroup our ‘phylogenomic variant set’. Our ‘phylogenomic variant set’ contained 94,993 biallelic SNPs, compared to 89,045 variants for our ‘population genomic variant set’.

To understand population structure and contemporary patterns of hybridisation, we inferred ancestry proportions for samples across all 16 populations using the likelihood-model approach in ADMIXTURE (Alexander *et al*., 2009), with *K* genetic clusters ranging from 1 to. We used all 89,045 SNPs in our filtered dataset described above. We converted data from VCF format to PLINK format using PLINK (v1.90b) and ran 3 iterations of ADMIXTURE for models up to *K* = 4. We evaluated the fit of the four models using the cross-validation procedure.

### Demographic modelling and inference of gene flow

To estimate the prevalence of introgression between *T. commodus* and *T. oceanicus* and the spatial context of gene flow, we estimated Patterson’s *D* statistic in *Dsuite* (Malinsky *et al*., 2020) across 94,993 biallelic SNPs. Specifically, we calculated D and f4 statistics to estimate gene flow between *T. oceanicus* (P2) and *T. commodus* (P3) by grouping populations by species and using *T. occipitalis* as an outgroup and *T. marini* (P1) as a derived species. Separately, we assessed D and f4 statistics across all possible combinations of populations in trios resulting in 1,141 comparisons of three different site patterns (ABBA, BBAA, BABA). Standard jackknife procedure across blocks was used to assess significance, with P-values corrected using Benjamini & Hochberg false-discovery rate correction (Benjamini & Hochberg, 1995). *D* statistics were considered significant only when Z-scores were larger than 3, remained significant following multiple correction (p < 0.05), and where *D* was larger than 0.05.

To accompany estimates of gene flow, we inferred the most likely demographic history of speciation between *T. commodus* and *T. oceanicus* using *dadi* (Gutenkunst *et al*., 2009) and a modular *dadi* pipeline (Portik *et al*., 2017). We converted our filtered VCF containing 89,045 biallelic SNPs to a joint site-frequency spectrum using easySFS (https://github.com/isaacovercast/easySFS). The SFS was projected down to 168, 242 for *T. oceanicus* and *T. commodus*, respectively. Consecutive optimisation rounds were performed for each model, with 4 multiple replicates (replicates = 10, 20, 30, 40) in each round. The best scoring parameter estimates (based on log-likelihood) were used to seed searches in the following round. Parameters were optimised using the Nelder-Mead method.

We first considered a series of simple demographic scenarios: (1) a ‘no migration’ model where species divergence occurred with no gene flow (included three parameters: divergence time, N_e_ for species A, N_e_ for species B), (2) a ‘symmetric migration’ model where species divergence was followed by a continuous period of symmetric gene flow (included a fourth parameter of constant gene flow), (3) an ‘ancestral migration’ model where divergence occurred with gene flow and ceased at some time point (included a fifth parameter of a second time denoting the end of gene flow), and (4) a ‘secondary contact’ model where symmetric migration occurs after some timepoint (included a fifth parameter of a second time denoting the beginning of gene flow).

Additional, more complicated, models were also considered: (5) an asymmetric migration model where species divergence occurred with a continuous period of asymmetric migration (included 5 parameters: divergence time, N_e_ for species A, N_e_ for species B, migration from species A to species B, migration from species B to species A), (6) an asymmetric migration model with an instantaneous size change parameter where species diverged with a continuous period of asymmetric migration and at some timepoint N_e_ changes in both species (included two additional parameters: N_e_ after size changes in species A, and N_e_ after size changes in species B), (7) an ancient migration model with asymmetric migration, (8) an ancient migration with asymmetric migration and an instantaneous population size change in both species, and (9) a secondary contact model with asymmetric migration and an instantaneous population size change in both species. Supplementary Figure 1 shows models and describes parameters. Finally, we scaled parameter estimates by first calculating N_ref_ = theta ÷ (4 x μ x effective sequence length), utilizing the *D. melanogaster* mutation rate μ (2.8 x 10^-9^) (Keightley *et al*., 2014), and where effective sequence length was the final number of SNPs included in the analysis.

Finally, we tested the hypothesis that the origin of *T. marini* may be a case of homoploid hybrid speciation with contributing ancestry from *T. commodus* and *T. oceanicus,* by calculating the f3 statistic in Treemix using *T. marini* individuals and populations of *T. commodus* and *T. oceanicus*. We converted our filtered VCF containing 89,045 biallelic SNPs to Treemix format using the Pop-gen pipeline platform (Webb *et al*., 2021). Standard errors for the f3 test statistic were calculated by jackknife procedure using blocks of 50 SNPs.

### Population genetic summary statistics

We calculated population genetic summary statistics across the genome to understand genetic differentiation and divergence between allopatric and sympatric populations of *T. oceanicus* and *T. commodus.* We first estimated site-wise Weir and Cockerham’s FST using VCFtools (v0.1.16)(Danecek *et al*., 2011) for five different comparisons using subsets of our filtered SNP dataset: (1) between all *T. commodus* and *T. oceanicus* individuals, (2) between only allopatric populations of *T. commodus* and *T. oceanicus* individuals, (3) between only sympatric populations of *T. commodus* and *T. oceanicus* individuals, (4) between allopatric and sympatric populations of *T. commodus* and (5) between allopatric and sympatric populations of *T. oceanicus*.

Separately, we calculated between-species FST, sequence divergence (d_XY_) and nucleotide diversity (π) by first re-calling variants alongside invariant sites. We filtered sites in the VCF as described above (minimum depth (10), max missingness (0.5), minimum quality score (30), minor allele count (3) and individuals with missing genotypes at half of all variable sites were removed). We estimated d_XY_ and π using Pixy (Korunes & Samuk, 2021) in 10kb windows with 10kb step sizes. We calculated d_XY_ between *T. commodus* and *T. oceanicus* individuals, and d_XY_ between both species in sympatry and allopatry separately. Functional identification was performed using custom scripts which primarily included blasting coding sequences against UniProt databases. Gene ontology for between-species F_ST_ outliers was performed in STRING <u>(Szklarczyk *et al*., 2015)</u>.

### Genetic associations for CHC, female preference, and male calling song traits

We performed genotype-phenotype association tests for three sexually selected traits – male calling song, intraspecific female calling song preference, and cuticular hydrocarbon pheromone profiles – involved in species discrimination. These three phenotypes were scored in *Teleogryllus oceanicus* only. This precluded us from testing genetic associations with traits directly involved in interspecific mating discrimination; however, it enabled us to test whether loci implicated in intraspecific mating variation show enhanced divergence in targeted interspecific comparisons, providing a powerful and unusual test of the connection between intraspecific sexual selection and interspecific divergence. We used RAD-seq data for F_3_ offspring previously produced from two mapping families described in Pascoal *et al*. (2020). Families were set up by initially crossing a sire from a Kauai, Hawaii stock line and a virgin dam from a Daintree, Australia stock line. These had been selected to maximize intraspecific variation in sexually selected traits of interest (Pascoal *et al*., 2017). Subsequently, F1 full-sib matings and then F2 full-sib matings were performed to produce 10 F3 mapping families consisting of 192 females and 199 males (of which 113 had ‘normal-wing’ venation which produces audible acoustic signals, and 86 had silent ‘flatwing’ venation owing to an adaptive male-silencing trait that is polymorphic in Hawaii) (Zuk *et al*., 2006; Pascoal *et al*., 2014, 2020).

#### Male calling song phenotyping

Following Moran et al. (2020), we scored 14 parameters of *T. oceanicus* male calling song in the F3 mapping population, which we considered as independent, continuous phenotypic variables. The parameters are can be observed in Supplementary Table 2 and all but carrier frequency are related to duration and temporal patterning of chirps within the standard, 2-component *T. oceanicus* song phrase. It is important to note that all three *Teleogryllus* species have similar 2-component songs consisting of a higher-amplitude trill followed by lower-amplitude chirps, but with different frequency and temporal features, lending plausibility to the hypothesis that loci underlying intraspecific song variation in *T. oceanicus* might also be under selection during interspecific encounters. Male calling songs were recorded using a Sennheiser ME66 microphone under red light between 22-25 °C (with the exception of a single individual recorded at 19 °C). For each individual, 5 song phrases from a continuous recording of male calling song were manually analysed using Sony SoundForge (v.7.0a). Each parameter was then averaged within each individual to obtain a set of 14 calling song traits for association mapping. We used song data from 84 mapping individuals after retaining only those that were successfully phenotyped as well as genotyped.

#### Female preference phenotyping

Females evaluate and respond to singing males by moving towards their song using directional phonotaxis. To score intraspecific female preferences, we exposed virgin females between the ages of 8 – 13 days to two male calling songs simultaneously and quantified their choice. We performed four successive trials for each female, with a minimum of 24 hours and maximum of 48 between trials. Assays were performed in a 172 cm x 28 cm arena lined with acoustic dampening foam. Population-average synthetic *T. oceanicus* song models corresponding to the original source populations of the crossing parents – Daintree in Queensland, Australia, and Wailua in Kauai, Hawaii – were broadcast at 70 dB at the female’s starting point using Sony SRS-m30 speakers mounted on either end of the arena. Average song parameters were calculated following the procedure in the above section, but for calling males from laboratory stock lines originally derived from the parental populations in Australia (*N* = 19) and Hawaii (*N* = 24) and with 10 song phrases measured per individual. We used two separate calling song playbacks for each species, which we constructed by artificially excising sound pulses from the laboratory recordings and using Sony SoundForge (v.7.0a) to assemble these into a temporal pattern fitting population-average chirp and interval durations and matching the necessary carrier frequency. Playback song parameters are given in Supplementary Table 2. We alternated the speaker direction and combination of song models between successive trials to avoid novelty and directional response biases. Thus, females were permitted to repeatedly choose between average male calling songs representative of either population. A positive response was scored if the subject made physical contact with a speaker within 5 minutes, and in that case the response latency was recorded. Before each test, females were given an acclimation period of 2 minutes underneath an inverted 100 mL plastic deli cup, and between each test the surface of the arena on which crickets walked was wiped down with 70% EtOH to minimize detection of odor cues. We used female choice and latency data from 59 mapping individuals after retaining only those that were successfully phenotyped as well as genotyped.

#### Cuticular hydrocarbon phenotyping

CHC profiles for mapping individuals were obtained from previously published data in Pascoal et al. (2020) and methods are fully described in Pascoal *et al*. (2016, 2020). To summarise, individually flash-frozen *T. oceanicus* crickets were immersed in HPLC-grade hexane (Fisher Scientific) for 5 minutes to remove long-chain waxy hydrocarbons that coat the insect cuticle. The resulting material was analysed with an internal pentadecane standard on an Agilent 7890 gas chromatograph linked to an Agilent 5975B mass spectrometer. Instrument run conditions were previously optimized for *T. oceanicus* (Pascoal *et al*., 2016). Mass spectrometry was performed using a C_7_-C_40_ alkane standard to enable calculation of peak retention indices (Supplementary table 3). For each sample, 26 peaks were integrated and quantified using MSD CHEMSTATION (v.E.02.00.493). The resulting peak abundances were entered into a principal components analysis (PCA) to reduce dimensionality for association mapping. Both sexes including two polymorphic male forms (silent ‘flatwing’ and singing ‘normal-wing’) present in the Hawaiian mapping population were combined in the PCA because we were primarily interested in the genetic architecture and loci associated with major sources of intraspecific CHC variation irrespective of sex or morph, as CHCs of both sexes are subject to sexual selection in this species (Thomas & Simmons, 2009, 2010). Accounting for individuals that were successfully phenotyped as well as genotyped, PCs 1-4 of 328 individuals entered the association mapping pipeline described below.

#### Association analyses

We conducted genotype-phenotype association analyses using a Wald test in GEMMA (v0.98.3)(Zhou & Stephens, 2012, 2014). For this, we fitted univariate linear mixed models to test for the association between each song parameter and SNP genotypes, accounting for population structure by jointly incorporating an inter-individual relatedness matrix along with individual mass and individual pronotum lengths as covariates. For CHCs, we fit a multivariate linear mixed model to test for an association between genetic variation and all four PC axes capturing most CHC variation (63.15%) among individuals. More PC axes could not be included the multivariate model due to computational limitations in GEMMA. In both analyses, we aimed to identify loci associated with intraspecific variation in song and CHCs, and variation that might be important in maintaining isolation between *T. oceanicus* and *T. commodus*. To assess significance, we adjusted P-values using a Bonferroni-Hochberg procedure with a significance threshold of corrected *P* < 0.05. To visualize these results, we plotted the log-transformed corrected p-values and switched their sign. Finally, we tested whether significantly associated song and CHC SNPs are implicated in divergence between *T. oceanicus* and *T. commodus* by first assessing the overlap between 99^th^ quantile F_ST_ and d_XY_ 10 Kb windows in our interspecific genome outlier scans with song and CHC-associated SNPs. Subsequently, we compared mean F_ST_ and d_XY_ 10 Kb windows for song and CHC associated regions with genome wide mean F_ST_ and d_XY._ We tested differences in means using two-sample t-tests.

We calculated the integrated haplotype score (iHS) to test for evidence of selective sweeps in two candidate regions associated with short-chirp inter-chirp intervals. We first phased our filtered SNP dataset using SHAPEIT4 with default parameters (Delaneau *et al*., 2019). We then used the R package *rehh* to filter data for minor allele frequency (minimum MAF<0.05) and run iHS scans for *T. commodus* and *T. oceanicus* separately. To identify genes implicated in song divergence, we looked for genes showing extreme iHS values (99^th^ percentile) and containing a SNP with significant association to a song parameter (particularly, the short-chirp inter-chirp interval parameter) using the *bedtools intersect* function.

#### Genotype-environment associations

To infer whether mating traits may potentially be under ecological selection or if local adaptation across environmental gradients confound signals of divergent selection between *T. commodus* and *T. oceanicus*, we performed genotype-environment association (GEA) scans. GEA approaches work by testing whether SNP allele frequencies are significantly associated with relevant environmental variables, after correcting for population structure. Here, since *T. oceanicus* and *T. commodus* populations show a clear latitudinal gradient, we utilize the latitude and longitude of each population as environmental variables. Latent factor mixed models are univariate models which test the association between each SNP and an environmental variable, and are among the most powerful methods to detect genotype-environment associations (Forester *et al*., 2018). These approaches require estimation of background structure via inference of the number of latent factors (*K*) in the dataset. We used sparse nonnegative matrix factorization (snmf) to estimate the number of latent factors and chose an optimal *K* (*K*=6) based on which *K* had the lowest cross-entropy criterion score, using the *R* package LEA (Frichot *et al*., 2014; Frichot & François, 2015).

LFMM was performed using a ridge regression on the full set of 89,045 biallelic SNPs and parameterized using a K=6, *via* the *R* package *lfmm* (Frichot *et al*., 2013; Caye *et al*., 2019)*. P*-values were calibrated for each SNP using a genomic inflation score. We conservatively characterised loci as “environmentally-associated” if calibrated p-values for both latitudinal and longitudinal association tests were *P* < 0.001. We tested the overlap between genes containing environmentally-associated SNPs and genes containing song and CHC-associated SNPs.

## Results

### Population structure

Figure 2B shows genome-wide PCAs examining population genetic structure in *T. oceanicus* and *T. commodus*. The species were largely separated into two clusters based on PC1 (which explained 55.5% of total variation). Both species show comparable intra-species variation along PC axes. This analysis also included samples from the third sister species, *T. marini*, which co-occurs in North Queensland with *T. oceanicus* and differs from *T. oceanicus* and *T. commodus* in calling song and colouration (Otte and Alexander, 1983; Moran et al., 2020). *T. marini* samples show extreme values on PC3 (explaining 3.5% of total variance) and are distinct from T. *oceanicus*, with which it is sympatric, and *T. commodus*.

Clustering of samples by species identity in the PCA was supported by multispecies coalescent phylogenetic analyses. However, whilst populations of *T. commodus* were monophyletic, *T. oceanicus* populations were not (Figure 2C). Our phylogenetic inference also supports early divergence of *T. marini*, followed by *T. oceanicus* and then *T. commodus* populations, suggesting north-to-south evolutionary splitting along the geographical transect. We found no evidence for recent admixture between *T. oceanicus* and *T. commodus* in their contact zone, recapitulating findings from Moran *et al*. (2018b) (Figure 2D). Additionally, ancestry proportions under K=4 reveal a latitudinal gradient for *T. commodus*, but not in *T. oceanicus*.

### Demographic history and historic gene flow between *Teleogryllus* species

Whilst we found little evidence for contemporary gene flow between *T. commodus* and *T. oceanicus* in contact zone populations, we found significant levels of historical gene flow between both species (Figure 3A), and significantly elevated *D*-values in contact zone populations compared to allopatric populations (Figure 3C). The best-fit demographic models supported a scenario of historical gene flow in sympatry that continued after gene flow in allopatry stopped. Interestingly, patterns of gene flow were consistent for all population comparisons, but particularly consistent among *T. commodus* populations (Figure 3B), indicating ancient hybridisation events very early on in species divergence are likely responsible for correlated *D*-values observed between species, and sympatric *T. commodus* populations may have experienced more unidirectional gene flow from *T. oceanicus* than vice versa. This is consistent with two *T. commodus*-assigned individuals showing mixed ancestry in the contact zone (1B & 1D). To investigate this in more detail, we compared models of continuous, ancient and recent migration in our demographic modelling framework. We found best support for an ancient migration model with symmetric isolation based on consistently low AIC scores across optimization rounds and particularly in the final model optimization round, where model convergence is expected (Figures 3D and 3E). Finally, we tested for but ruled out a hybrid origin of *T. marini*, finding no evidence that it is admixed (Figure 3F).

**Figure 3:**
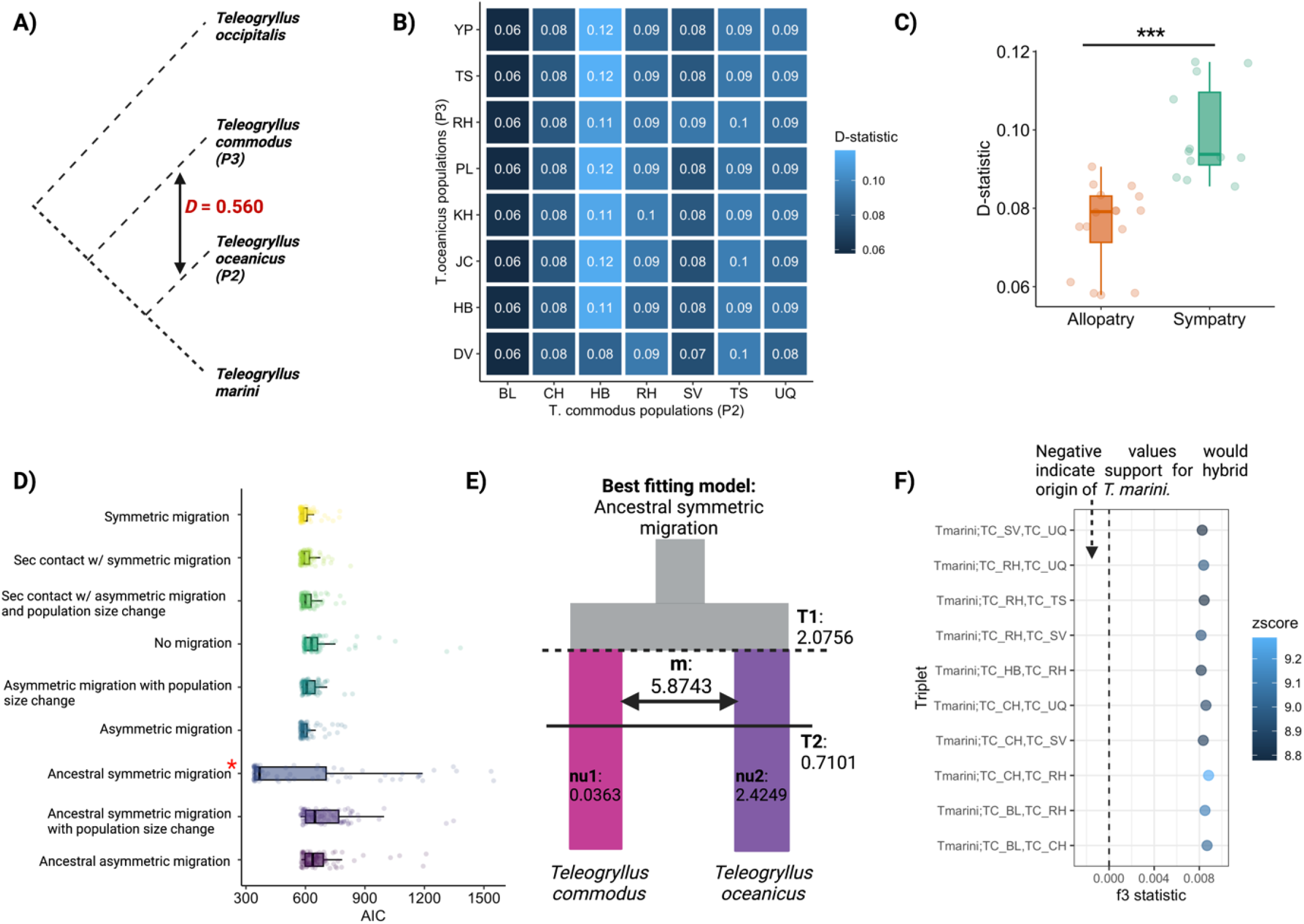
Population demography and historical gene flow. **A)** Inferred gene flow (*D* statistic) between *T. oceanicus* and *T. commodus.* **B)** Gene flow between *T. oceanicus* and *T. commodus* populations, with *T. occipitalis* as the outgroup. *D*-statistics were averaged for independent tests where P1 was permuted across all populations. Only comparisons with significant adjusted p-values after multiple testing and Z-scores > 3 were considered and shown here. **C)** Gene flow inference across allopatric and sympatric populations of *T. oceanicus* and *T. commodus.* Comparison of *D*-statistics for trios where P2 (*T. commodus*) and P3 (*T. oceanicus*) are allopatric or where both sister species are sympatric. Stars indicate level of significance: 0.0001 = ***, 0.001 = **, 0.01 = *, 0.05 = >0.05 = ns. **D)** Model fits for different species divergence scenarios with and without gene flow, using *dadi*. Each point shows AIC for an independent model-fitting run. Red asterisks shows best-itting model after 50 optimized, independent runs. **E)** Diagram showing the best-fitting model divergence with ancestral symmetric migration between *T. oceanicus* and *T. commodus*) unscaled parameter estimates of T1 (initial divergence parameter), T2 (time of cessation of gene flow period parameter), m (symmetric migration), nu1 (population size of *T. commodus*) and nu2 (population size of *T. oceanicus*). **F)** F3 test for hybrid origin of *T. marini.* Dashed line indicates zero; values below 0 with significant z-scores show support for hybrid origin for population C (C;A,B), where populations A and B represent parental lineages.

### Pronounced, asymmetrical genomic differentiation in sympatry

Average F_ST_ between species was 0.072, with significantly lower differentiation on the X chromosome (F_ST_ = 0.063) compared to the autosomes (F_ST_ = 0.074) (two sample *t-test*: *t* = 11.02, *p* < 0.0001) (Supplementary Figure 2). Average F_ST_ between *T. commodus* and *T. oceanicus* in sympatry (F_ST_ = 0.156) was considerably and significantly higher than in allopatry (F_ST_ = 0.088) (two sample *t-test*: *t* = −65.58, *p* < 0.0001) (Figure 4C and 4D). Heightened differentiation in the contact zone is attributable to greater divergence of *T. commodus* contact zone populations: average F_ST_ between sympatric *T. commodus* and allopatric *T. oceanicus* (F_ST_ = 0.155) replicated the considerable difference in average F_ST_ between sympatric populations of both species, but F_ST_ between sympatric populations of *T. oceanicus* compared to allopatric populations of *T. commodus* (F_ST_=0.0804) did not (see also Figure 4E and 4F compared to Figure 4G and 4H). Alongside evidence of stronger gene flow in sympatric populations, our results indicate genomic barriers to gene flow may be asymmetrically strengthened in the contact zone between *T. commodus* and *T. oceanicus*.

**Figure 4:**
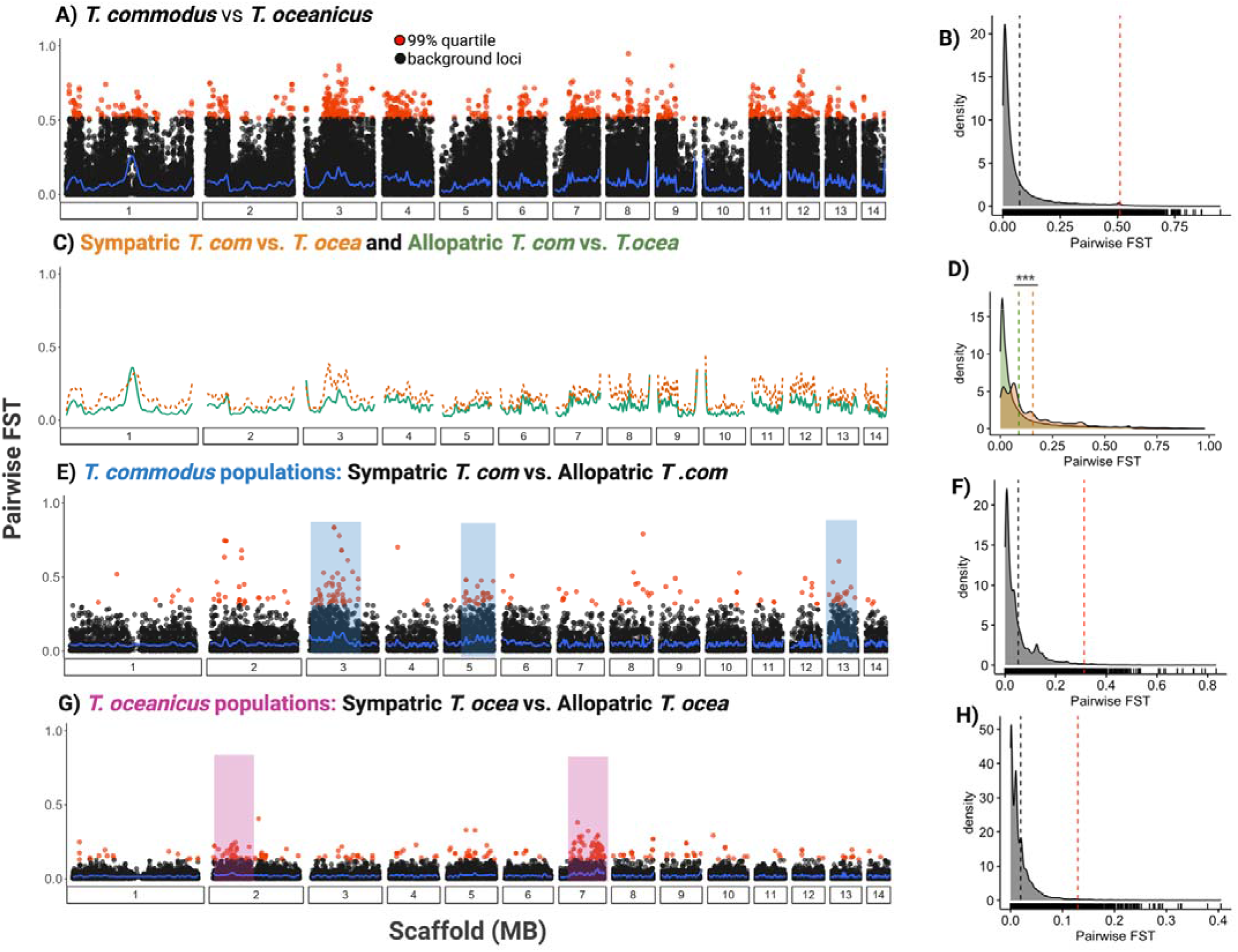
Population-pairwise F_ST_ between and within *T. commodus* and *T. oceanicus*. **A)** Site-wise between all *T. commodus* versus *T. oceanicus* population pairs. **B)** Density of sites (short vertical ck bars above x-axis), distribution of their pairwise F_ST_ (shaded region), mean F_ST_ (dashed black line) d 99^th^ quantile outlier mean (dashed red line). **C)** Smoothed site-wise F_ST_ comparing sympatric *T. mmodus* and *T. oceanicus* (dashed orange line) and allopatric *T. commodus* and *T. oceanicus* (solid en line). **D)** Distributions of pairwise F_ST_ in sympatric (orange) and allopatric (green) comparisons. Stars icate significance at p < 0.0001. **E)** Site-wise F_ST_ between *T. commodus* populations sympatric with *T. anicus* versus allopatric with *T. oceanicus.* Blue shading indicates genomic regions enriched for outlier i. **F)** Distribution of pairwise F_ST_ with mean (black dashed line) and mean for 99^th^ quantile outlier SNPs shed red line). **G)** Site-wise F_ST_ between *T. oceanicus* populations sympatric with *T. commodus* versus se allopatric with *T. commodus*. (H) Distribution of pairwise F_ST_ showing mean (black dashed line) and an for 99^th^ quantile outlier SNPs (dashed red line). Pink shading indicates genomic regions enriched for lier loci.

To identify genomic regions potentially involved in species divergence, we characterised F_ST_ outlier SNPs between *T. commodus* and *T. oceanicus*, and between sympatric and allopatric populations (intraspecies, interpopulation) within the same species for both species. We found 4,457 (99^th^ quantile) outlier SNPs between species out of 89,045 SNPs and identified 243 genes with functional annotations that overlapped with outlier SNPs. Apart from scaffold 10, outlier SNPs were distributed almost uniformly across the genome (Figure 4A and 4B). For the latter, a third of significant Gene Ontology terms relate to neural development (Supplementary Figure 3). To increase the robustness of inferences about potential genomic barriers to gene flow, we also calculated genome-wide d_XY_ in 10kb windows and identified 651 outlier windows (d_XY_ > 0.0149). Only 43 genes were found to overlap with the 651 d_XY_ outlier windows. We found 84 shared outlier regions and 17 shared genes when examining the intersection of F_ST_ and d_XY_ outliers. The latter included calmodulin-binding transcription factor (*CAMTA*), known to be involved in male courtship song in *D. melanogaster* (Sato *et al*., 2019) and associated with male song variation in *Lapaula* (Xu & Shaw, 2021).

To test the effect of sympatry on intraspecific variation, we identified intraspecies, interpopulation F_ST_ outliers by contrasting genetic differentiation between sympatric and allopatric populations for each species separately. Regions of increased differentiation were largely unshared between species (Figure 4E and 4G). For example, *T. commodus* contrasts identified 47 genes and *T. oceanicus* contrasts identified 83, but only 4 were shared (Supplementary Table 4).

### Loci associated with intraspecific song, female preference, and CHC variation

#### Male calling song

Five of the 14 male calling song traits showed significant genomic associations and in all cases these were distributed over more than one chromosome: long chirp pulse duration, the duration of long chirp-short chirp intervals, short-chirp pulse duration, short-chirp inter-chirp interval and inter-song interval (Figure 5A). There was negligible evidence for genomic co-localization of different song traits, highlighting the complex architecture of male calling song (see also Supplementary Figure 4) (Figure 5B). Overall, we found 162 significantly associated SNPs (corrected *P* < 0.05), and these overlapped or were within 10 kb of 179 genes. These included orthologs in *D. melanogaster* that modulate courtship behaviour (*takeout* and *lola*) (Dauwalder *et al*., 2002; Lazareva *et al*., 2007; Dinges *et al*., 2017; Sato *et al*., 2019).

**Figure 5.**
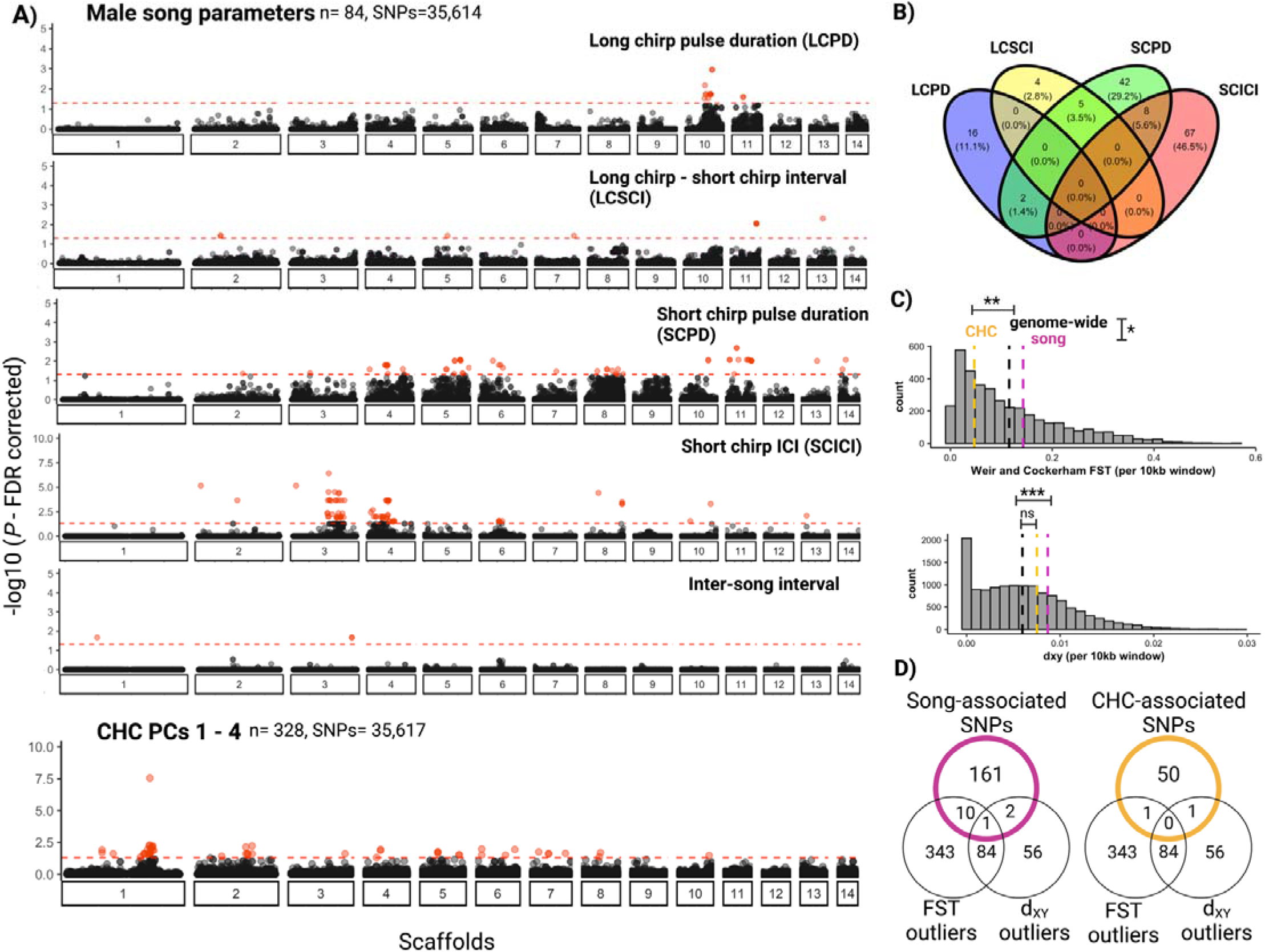
Linking intraspecific genetic variation for male calling song and CHCs with rspecific barrier loci in *Teleogryllus* spp. **A)** Significant genome-wide association (GWA) yses. Only song parameters with at least one significant association are shown. Bottom panel ws GWA analysis for CHCs, combining sexes. Dotted red line and points indicate significance shold (FDR-corrected P < 0.05) and associated SNPs, respectively. **B)** Venn diagram showing lap of male song trait SNPs. **C)** Histograms of F_ST_ and d_XY_ calculated in 10 kb windows. Dotted and orange lines indicates mean F_ST_ and d_XY_ for 10 kb regions containing at least one SNP stically associated with song and CHC variation, respectively. The dotted black line indicates ome wide mean F_ST_ and d_XY._ Stars indicate level of significance for two-sample t-tests: 0.0001 *, 0.001 = **, 0.01 = *, 0.05 =., >0.05 = ns. **D)** Overlap of genes within F_ST_ and d_XY_ outlier ons (99^th^ quantile of distributions), and song and CHC-associated SNPs.

#### Cuticular hydrocarbons

Altogether, we found 50 SNPs (Wald test, corrected *P* < 0.05) associated with variation in CHC production across 328 individuals. These SNPs overlapped or were within 10kb of 22 genes (Supplementary Table 5), but none of the genes have previously been implicated in CHC variation or production. Consistent with previous analyses (Pascoal *et al*., 2020), we find a large peak of association for CHC production on the X chromosome (scaffold 1). Of the 50 significantly-associated SNPs, 20 span a 4.2 MB region on the X chromosome containing *doublesex,* a candidate region implicated in a male wing-feminising adaptation to an acoustically-orienting parasite in Hawaiian populations of *T. oceanicus* (Figure 5A; bottom panel) (Zhang *et al*., 2021). When searching for sex-specific effects of CHC production, we found no associations after correcting for multiple testing.

#### Female preference

We found no significant genomic associations for female preference for male calling song, or latency of female phonotaxis response to calling song, after correcting for multiple testing (Supplementary Figure 5). Female mate choice was therefore not considered in subsequent analyses.

#### Genotype-environment association tests

We identified 867 SNPs (P < 0.001) associated with latitudinal and longitudinal gradients across the 16 populations sampled. These SNPs corresponded to 184 genes with annotations. We asked whether any of the song or CHC-associated genes showed allele frequency changes associated with latitudinal and longitudinal variables. We found only four environment-associated genes which also showed significant song-associations (out of 179 song-associated genes). These genes include *seven-up* and *methuselah-like 2* (mthl2). Similarly, only two CHC-associated (out of 22) genes showed significant associations with both latitudinal and longitudinal variables.

### Association of intraspecific sexual trait loci with interspecific divergence

To test for relative importance of genes involved in intraspecific male courtship song and CHC variation in interspecific divergence, we compared F_ST_ and d_XY_ in 10kb windows containing SNPs (and presumably gene regions) associated with song and CHC variation with background levels of genomic divergence between *T. oceanicus* and *T. commodus*. Whilst song genes show significantly higher genomic differentiation and divergence compared to the genetic background (two-sample t-tests: F_ST_, *t* = −2.16, *P* = 0.036; d_XY_, *t* = −4.31, *P* < 0.0001), genes associated with intraspecific CHC variation showed significantly lower differentiation, but comparable genomic divergence to the genetic background (two-sample t-test: F_ST_, *t* = 3.15,*P* = 0.005; d_XY,_ *t* = −1.87, *P* = 0.06) (Figure 4C). Additionally, we tested for evidence of stronger diversifying selection of song and CHC-associated genes in sympatry compared to allopatry and found no difference in F_ST_ between song-associated (Wilcox test; W = 1712, *p* = 0.9) and CHC-associated (Wilcox test; W = 279, *p* = 0.3) genes in sympatry compared to allopatry.

For CHC variation, one gene showed signal of high genomic differentiation (*phtf*) and one gene showed signal of high genomic divergence (*RYK*), suggesting genes associated with intraspecific CHC variation are not involved in interspecific divergence. However, we found 10 intraspecific song-associated genes with high genetic differentiation between *T. commodus* and *T. oceanicus*, and 2 genes (*semphaorin-2A* and *Syntrophin-like 2)* showing signals of high genomic divergence. The gene *semaphorin-2A* (SEM2A) is significantly associated with short-chirp inter-chirp interval (Scaffold 4, position: 44,431,198; Wald test, uncorrected *P* < 0.0001 and corrected *P* = 0.01) and shows elevated signals of differentiation (F_ST_= 0.53) and divergence (d_XY_= 0.02). Overall, we found no evidence that song-associated genes were over-represented in regions of high genomic differentiation (Fisher’s exact test; *P* = 0.3).

To test if song genes were under recent positive selection in *T. commodus* and *T. oceanicus*, we focused on scaffolds 3 and 4, where large genomic regions show consistent, significant associations with the short-chirp inter-chirp interval. Within those focused regions, and globally across scaffolds 3 and 4, we calculated species-specific integrated haplotype scores (iHS) and asked which song-associated genes, if any, showed signatures of selective sweeps. Examining the most extreme values of the haplotype statistic (iHS – 99^th^ percentile), we found no scaffold 3 genes that met our conservative iHS cut-off and were significantly associated with any song parameter in *T. commodus*, and only one gene (*Tudor-SN*) associated with short-chirp inter-chirp intervals and showed evidence of a selective sweep in *T. oceanicus* (iHS = 3.31, log *P =* 3.03). For scaffold 4, we similarly found no overlap between iHS outliers and song-associated genes in *T. commodus*, but we found 6 genes that met both criteria in *T. oceanicus*, including *semaphorin 2A*, *seven-up,* and *Syntrophin-like 2*.

## Discussion

Critical tests that link the genetics of intraspecific variation in mating traits with the genetics of reproductive isolation between species are rare, despite their importance for testing seminal models of speciation by sexual selection (Kopp *et al*., 2017). Our analysis of Australian field cricket sister species provides a rare opportunity to support or refute predictions about this widely-discussed but poorly understood link between microevolutionary process and macroevolutionary pattern: overall, there was limited evidence that genes underlying sexual traits within *T. oceanicus* are generally implicated as candidate barrier loci between *T. oceanicus* and *T. commodus*; yet those that are show evidence of divergent selection that is not clearly attributable to ecological selection. We discuss the implications and limitations of these findings and identify areas where specific experimental advances will be required to overcome common obstacles that impede understanding of the genomics of speciation via sexual selection.

Our study demonstrates that *T. oceanicus* and *T. commodus* are genetically distinct and show little evidence of contemporary hybridization. Nevertheless, these sister species show striking patterns of heightened genome-wide differentiation in regions of sympatry despite evidence of elevated levels of historical gene flow in sympatry compared to allopatry. Such genomic signals are consistent with reproductive character displacement in sympatry (Garner *et al*., 2018), suggesting a role for reinforcement in driving divergence between *T. oceanicus* and *T. commodus*. This increase in genomic differentiation in sympatry may be the result of asymmetric strengthening of barriers to gene flow in sympatric *T. commodus* populations. Loci associated with intraspecific variation in male advertisement song and CHC profiles did not collectively show evidence of divergent selection between species, but the small number that did may play a disproportionate role in species divergence. Intriguingly, the latter were mostly limited to loci associated with advertisement song variation within *T. oceanicus* but less so CHCs, suggesting co-option of the genetic architecture of intraspecific mating signals depends on the presence or absence of constraints imposed by countervailing sources of selection.

So far, only a few studies have investigated whether genome-wide patterns of differentiation reflect a signal of reinforcement (Hopkins *et al*., 2012; Smadja *et al*., 2015). We found evidence of significant, increased differentiation, owing to divergence in sympatric *T. commodus* populations. This genomic analysis is consistent with the greater phenotypic divergence of male advertisement song and CHC profiles observed in the *T. oceanicus* and *T. commodus* contact zone (Moran *et al*., 2020). Distinguishing between scenarios that may explain the patterns observed here is challenging. For instance, whilst we have shown evidence of uniform increase in genetic differentiation in sympatry mirroring concomitant evidence of character displacement in the contact zone between *T. oceanicus* and *T. commodus*, it is unclear whether it is (solely) reinforcement driving this signal (Noor, 1999; Hoskin & Higgie, 2010). Both *T. oceanicus* and *T. commodus* are collected in the same localities and occupy similar ecological niche space, suggesting local adaptation and ecological character displacement do not best explain the patterns observed here. Using genotype-environment association tests, we demonstrate that there is little overlap between genomic loci associated with sexual traits and genes potentially under local adaptation, indicating little support for a role of ecology in driving divergence of mating traits.

For the 5 song parameters that showed significant statistical associations, we show that their genetic bases are largely polygenic and do not co-localize (Figure 4A & B), consistent with expectations of a complex genetic basis of male courtship song in other insects such as *Drosophila* (Gleason & Ritchie, 2004; Turner *et al*., 2013; Pischedda *et al*., 2014). Candidate barrier loci were dominated by genes enriched for neurological functions, indicating potential behavioural rewiring, and several of the small number of genes enriched in both intraspecific and interspecific analyses have been implicated in the control of courtship song variation in other species (Turner *et al*., 2013; Blankers *et al*., 2018; Sato *et al*., 2019; Xu & Shaw, 2019). For example, genes identified in our outlier scans may also play a role in species divergence in a rapid radiation of endemic Hawaiian crickets, and several have been reliably annotated as *D. melanogaster* orthologs and were previously implicated in *Drosophila* courtship behaviour (Supplementary Table 6). *Lola* is one such promising candidate. In *Drosophila*, it is involved in neuronal reprogramming that interacts with *fruitless*, a regulator of courtship circuitry; it is associated with long-chirp pulse duration in *T. oceanicus* in this study (corrected *p*: 0.018); and it is also under a moderate-effect QTL for mating song rhythm variation in *Laupala* (Dinges *et al*., 2017; Blankers *et al*., 2018; Sato *et al*., 2019). Some of the song-associated genes found in our genome-wide association tests were also outliers in our F_ST_ and d_XY_ genome scans and represent additional candidates for further study. For example, *Syn2* is associated with variation in short-chirp inter-chirp intervals and shows evidence of positive selection in this study via extended haplotype homozygosity tests, and *Syn1* in *Drosophila melanogaster* has been associated with variation in inter-pulse intervals of courtship song through an Evolve and Resequence experiment (Turner *et al*., 2013).

Our findings contrast with other genomic studies of courtship song and preference which suggest that courtship song is often underpinned by large effect loci (Ding *et al*., 2016), and that sexual signals and preference are underpinned by co-localized gene clusters (Shaw & Lesnick, 2009; Xu & Shaw, 2019, 2021). Because we found a multitude of loci associated with male advertisement song parameters in *T. oceanicus*, rapid divergence of multiple song parameters in *T. oceanicus* and *T. commodus* probably requires polygenic selection, which may be considerably more difficult to evolve than if calling song was underpinned by simpler genetic architecture (Kopp *et al*., 2017). We found no statistically significant associations for female preference, likely due to difficulty in accurately capturing and mapping a transient behavioural decision and, in this case, limited by sample size (n=59), so we could not assess whether genes associated with male advertisement song are also associated with female preference. We recapitulate findings from Pascoal *et al*., (2020) and show that close to half the associated SNPs found in our genome-wide association for CHC production are found in a 20 Mb region around *doublesex* (Pascoal *et al*., 2014, 2020). However, only two SNPs associated with CHC production are found within or around genes that are also outliers in our genome scans, and genes associated with intraspecific CHC production are significantly less differentiated compared to the genomic background (Figure 5C), suggesting they may be under strong selective constraint between species. In contrast with song-associated genes, patterns of genetic variation in CHC-associated genes and underlying phenotypic variation may be mediated by natural and sexual selection acting in opposing directions, which is a central prediction for classic models of sexual selection <u>(Fisher, 1915, 1958; Lande, 1981; Kirkpatrick, 1982</u>; <u>see also Berson *et al*., 2019)</u>.

### Conclusions

The evolution of divergent mating traits is an important mechanism that can restrict gene flow between nascent lineages and maintain species boundaries during species coexistence. Our results suggest that the genetic architecture of intraspecific variation in sexual traits can be co-opted during species divergence, but that this co-option is subject to important limitations. In *T. oceanicus*, such co-option is likely driven by polygenic selection of a limited subset of loci controlling acoustic signaling. Chemical signals are similarly polygenic, but under considerable ecological constraint and therefore not as important for divergence between *T. commodus* and *T. oceanicus*. Our study represents one of few dual assessments of the genetic basis of intraspecific mating traits and divergent interspecific selection in natural populations. Our findings support a scenario where few, but essential, loci play a dual role in intraspecific sexual selection and interspecific divergence, whereas many other loci contribute to either process but do not mechanistically link the two. The quest to connect microevolutionary process and macroevolutionary pattern during speciation by sexual selection might therefore profit from combinatorial approaches such as we present here to narrow the focus on rare ‘speciation-and-sexual-selection’ genes.

## Author contributions

L.H.Y: conceptualization, data curation, formal analysis, investigation, methodology, visualization, writing—original draft, and writing – review and editing. S.P: data curation, investigation, methodology, resources. PAM: data curation, methodology, resources, and writing – review. NWB: conceptualization, funding acquisition, investigation, methodology, resources, writing—original draft, and writing—review & editing.

## Acknowledgements

We thank J.G. Rayner for useful discussion and feedback in preparing the manuscript, and gratefully UK Natural Environment Research Council for funding support. We also thank the Crop Diversity High Performance Cluster (HPC) for bioinformatic assistance. The authors declare no conflicts of interest.

## Data accessibility

Raw sequence data used in this paper can be found in NCBI under the BioProject IDs: PRJEB26502 (population genomics) and PRJEB24786 (genotype-phenotype mapping). Male calling song, female preference and CHC data will be available as supplementary material and archived in Dryad.

